# Cooption of polyalanine tract into a repressor domain in the mammalian transcription factor HoxA11

**DOI:** 10.1101/2020.02.19.956243

**Authors:** Vincent J. Lynch, Gunter P. Wagner

## Abstract

An enduring problem in biology is explaining how the functions of genes originated and how those functions diverge between species. Despite detailed studies on the functional evolution of a few proteins, the molecular mechanisms by which protein functions have evolved are almost entirely unknown. Here we show that a polyalanine tract in the homeodomain transcription factor HoxA11 arose in the stem-lineage of mammals and functions as an autonomous repressor module by physically interacting with the PAH domains of SIN3 proteins. These results suggest that long polyalanine tracts, which are common in transcription factors and often associated with disease, may generally function as repressor domains and can contribute to the diversification of transcription factor functions despite the deleterious consequences of polyalanine tract expansion.

**Research Highlights:** We show that a polyalanine track in HoxA11 evolved into a repressor domain in mammals through an increase in alanine repeat number, indicating that transcription factors can evolve novel functions despite the potential deleterious consequences associated with amino acid repeats.

## Introduction

The mechanisms of gene regulatory evolution are debated, however, it is clear that changes in the regulatory activities of transcription factors plays an important role in the evolution of gene regulation (Wilson, 1975). Detailed functional dissection of a few transcription factors indicates that the evolution of new co-factor interactions (Löhr et al., 2001; Heffer et al., 2010; Brayer et al., 2011b), ligand-binding activities (Thornton et al., 2003; Ortlund et al., 2007; Bridgham et al., 2009), subcellular localization signals (Salichs et al., 2009), post-translational modifications (Lynch et al., 2011), and neo-allosteric effects (Nnamani et al., 2016) can generate functional diversity. Despite these studies, major questions in transcription factor evolution remain unanswered. For example, how have transcription factors evolved and modularized their regulatory functions, how are these functions maintained despite sequence divergence, and how do functional changes occur without strong negative pleiotropic effects?

Simple sequence repeats, such as long tracts of the same amino acid (homopolymers), are common in eukaryotic transcription factors, particularly developmental genes (Karlin and Burge, 1996; Briata et al., 1997; Albà et al., 1999; Young et al., 2000; Janody et al., 2001; Albà and Guigo, 2004; Faux et al., 2005). These regions are extremely variable between species (Briata et al., 1997; Albà et al., 1999; Albà and Guigo, 2004) and have been associated with morphological evolution (Galant and Carroll, 2002; Ronshaugen et al., 2002; Fondon and Garner, 2004; Anan et al., 2007), suggesting they contribute to the functional diversity of transcriptional regulators (King et al., 1997; Kashi and King, 2006; Lynch and Wagner, 2008). Expansions of these repeats also cause developmental defects and neurodegenerative disease (Karlin and Burge, 1996; Goodman et al., 1997; Albà and Guigo, 2004; Oma et al., 2007; Orr and Zoghbi, 2007). Polyalanine tracts, for example, account for ∼17% of all human homopolymeric protein sequences (Karlin et al., 2002) and expansions of polyalanine tracts in at least 11 genes have been associated developmental defects and disease (Brown and Brown, 2004). Despite their ubiquity and association with diseases, however, few functions have been identified for amino acid repeats (Emili et al., 1994; Gerber et al., 1994; Xiao and Jeang, 1998; Janody et al., 2001; Galant and Carroll, 2002; Salichs et al., 2009).

Alanine rich regions have long been recognized to be common in transcriptional repressor domains suggesting they have a role in mediating transcriptional repression (Janody et al., 2001; Maurer et al., 2003), but a mechanistic explanation for these observations has been lacking. Here we show that a polyalanine tract in the homeodomain transcription factor HoxA11 evolved in the stem-lineage of mammals, physically interacts with the PAH domains of SIN3 proteins, and functions as a SIN3A/HDAC1-dependent repressor domain. Remarkably, while our results suggest that polyalanine tracts may generally have the ability to function as SIN3-dependant repressor domains polyalanine tract expansions are also associated with several disease suggesting they can contribute to the functional diversification of transcription factors despite their potentially deleterious consequences.

## Materials and Methods

### Validation of IGFBP1 as a negatively regulated target gene of HoxA11

Human endometrial stromal fibroblasts (ESFs) immortalized with telomerase (ATCC, Cat. No. CRL-4003) were grown in steroid-depleted DMEM, supplemented with 5% charcoal-stripped calf-serum and 1% antibiotic/antimycotic (ABAM). For induction of decidualization, ESFs were treated with 0.5 mM 8-Br-cAMP (Sigma) and 1 µM medroxyprogesterone acetate (Sigma). At 80% confluency, cells were transfected with HoxA11 Silencer Select Validated siRNAs or siRNA controls (Ambion, Cat. 4390824) and Lipofectamine RNAiMAX (Invitrogen) according to the manufacturer’s protocols. Cells were harvested and RNA collected 48 hours after differentiation/transfection. Real-time PCR reagents, including TaqMan FAST Universal PCR Master Mix, and HoxA11, IGFBP1 and GADPH endogenous control primer/probe sets were purchased from Applied Biosystems.

### Ancestral sequence reconstruction

To determine when the polyalanine tract in HoxA11 evolved we identified *HoxA11* genes from 156 placental mammals, 3 marsupials, 2 monotremes, 8 sauropsids (including birds, turtles, and squamate reptiles), 2 species of amphibian, 2 species of coelacanth, 30 Euteleost fish, and 4 species of Chondrichthyes (**Fig. 2A**). Ancestral sequences were inferred with PAML (CODEML) using the species tree and the GTR model of sequence evolution, indels inferred by maximum parsimony. The Bayesian posterior probability at each site of the reconstructed ancestral sequence was >0.96 for all sites and genes. The ancestral Eutherian, Therian, Mammalian, and Amniote genes were synthesized by GeneScript Corp. with human optimized codon usage and ligated into pcDNA3.1(+)-V5/His as described previously (Brayer et al., 2011a; Nnamani et al., 2016). Proper expression and nuclear localization of all extant and reconstructed expression constructs were verified by western blotting. Interested readers are referred to (Brayer et al., 2011a; Nnamani et al., 2016) for further details.

**Figure 1.**
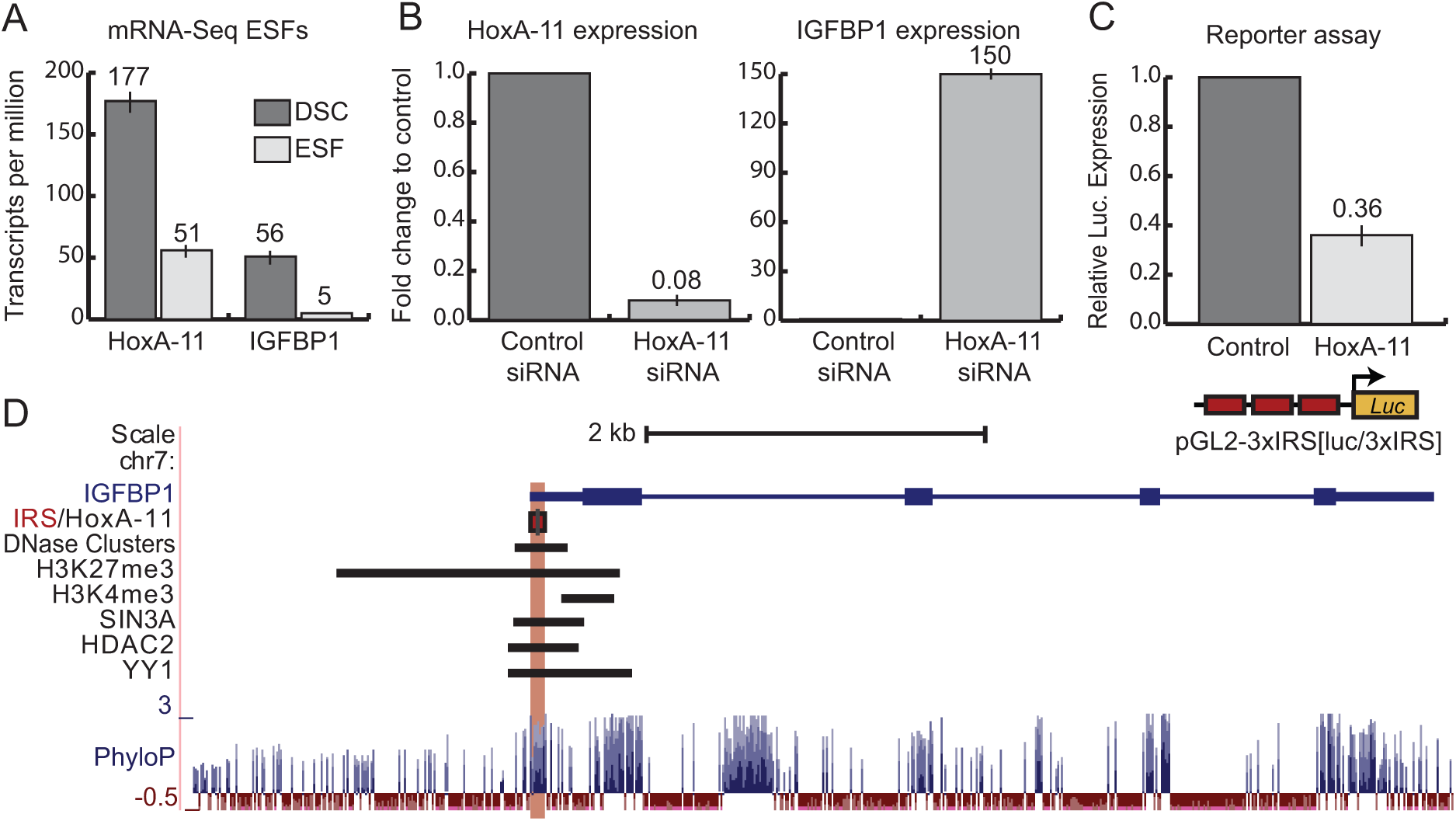
HoxA11 negatively regulates *IGFBP1* expression. **(A)** Expression of *HoxA11* and *IGFBP1* in mRNA-Seq data from human endometrial stromal fibroblasts (ESFs) and differentiated decidual stromal cells (DSCs). n=3, mean ± SD. **(B)** Expression of HoxA11 and IGFBP1 after siRNA-mediated knockdown of HoxA11 (n=3, mean ± SD). Results are shown relative to control siRNA and *OAS1* endogenous control. **(C)** HoxA11 represses luciferase expression from the pGL2-3xIRS[*luc*/3xIRS] reporter vector (n=4, mean ± SEM). **(D)** The location of the IRS is shown in red and the location of the Hox binding site as a black line. The location of DNase clusters, H3K27me3, SIN3A, HDAC2, and YY1 ChIP-Seq peaks are shown as black bars. Conservation of the region is shown as a PhyloP plot.

**Figure 2.**
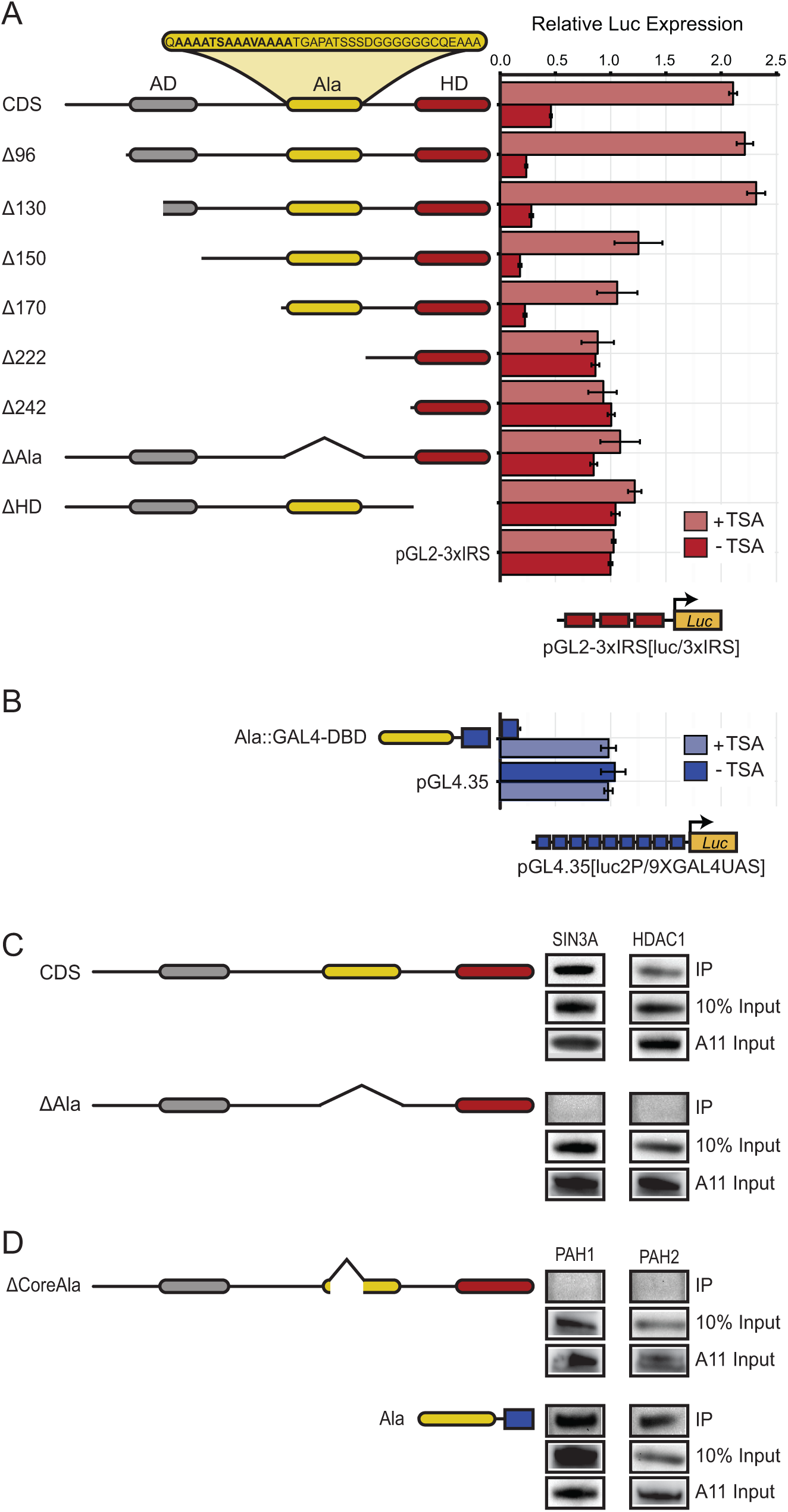
The HoxA11 polyalanine tract is a SIN3A/HDAC1-dependent transcriptional repressor domain. **(A)** Repression of luciferase expression from the pGL2-3xIRS[*luc*/3xIRS] reporter vector by full length (CDS) and deletion mutants (Δ) of mouse HoxA11 without (red bars) and with TSA (light red bars). Results are shown as fold changes in luciferase (Luc) expression relative to empty expression vector controls (n=4, mean ± SD). The relative location of the N-terminal activation domain (grey rounded rectangle, AD), the polyalanine tract (yellow rounded rectangle, Ala), and the homeodomain (red rounded rectangle, HD) are shown as cartoons illustrating the location of deletion mutants. The amino acid sequence of the mouse polyalanine tract is shown, with the core alanine repeat in bold) **(B)** Repression of luciferase expression from the pGL4.35[*luc2P*/9X*GAL4*UAS] reporter vector by the GAL4-DBD::polyalanine-tract fusion protein without (blue bars) and with TSA (light blue bars). Results are shown as fold changes in luciferase (Luc) expression relative to empty expression vector controls (n=4, mean ± SD). **(C)** Co-immunoprecipitation of the full length (CDS) and ΔAla mutant HoxA11 proteins with SIN3A and HDAC1. **(D)** Co-immunoprecipitation of the mouse HoxA11 core alanine deletion mutant (ΔCoreAla) and of GAL4-DBD tagged alanine tract with the V5/His-tagged PAH1 and PAH2 domains of SIN3A.

### Cell culture and luciferase reporter assays

*HoxA11* was amplified by PCR from embryonic cDNA from mouse (*Mus musculus*). Full-length and deletion coding regions were cloned into the GAL4 DNA-binding domain vector pM2 (mouse amino terminal and internal deletions), pBIND (alanine repeat), or pFLAG. HeLa cells were grown in DMEM supplemented with 10% FBS. Cells were transiently transfected with Lipofectamine 2000 (Invitrogen) according to the manufacturer’s protocol with 2ng of the *Renilla* control vector (pGL4.71) and 50ng of the appropriate luciferase reporter. Luciferase expression was assayed 48 hours after transfection using the Dual Luciferase Reporter System (Promega). Each experiment was repeated four times, with 8 replicates per experiment.

### Co-immunoprecipitation assays

HeLa cells were incubated overnight at a density of ∼4.4 × 10^6^ cells/mL on 10 cm plates prior to transfection. A total of 12 μg of appropriate expression vectors was transfected and incubated for 4 hours at 37°C, before addition of 7 mL DMEM. After 16 hours the transfection media was removed and replaced with fresh DMEM, and the cells were incubated an additional 24 hours before harvesting. After removing DMEM and washing cells twice with PBS, 1 mL ice-cold lysis buffer (20 mM Tris, pH 8.0, 40 mM KCl, 10 mM MgCl_2_, 10% glycerol, 1% Triton X-100, 1x Complete EDTA-free protease inhibitor cocktail (Roche), 1x PhosSTOP (Roche)) was added to each plate and cells were harvested by scraping with a rubber spatula. Cells were then incubated on ice for 30 minutes in 420 mM NaCl. Whole cell lysate was cleared by centrifugation at 10,000 rpm for 30 minutes at 4°C, and supernatant was transferred to a clean microfuge tube. After equilibrating protein concentrations, 1 mL of sample was mixed with 40 mL of antibody conjugated agarose beads (Sigma) pre-washed with TNT buffer (50 mM Tris-HCl, pH 7.5, 150 mM NaCl, 0.05% Triton X-100), and rotated overnight at 4°C. The following day, samples were treated with 50 U DNase (Roche) and 2.5 μg RNase (Roche) for 60 minutes at room temperature, as indicated. Samples were washed 3x with 1 mL wash buffer (150 mM NaCl, 0.5% Triton X-100). After the final wash, agarose beads were resuspended in elution buffer (500 mM Tris pH 7.5, 1 M NaCl), and boiled to elute immunoprecipitated complexes. Eluted protein was run on Bis-tris gels, probed with antibodies specific to epitope tags, and visualized by Chemi-luminescence.

## Results and Discussion

### HoxA11 is a transcriptional repressor of *IGFBP1*

Previous studies have found that the insulin response sequence (IRS) from the *insulin-like growth factor binding protein 1* (*IGFBP1*) enhancer is directly bound by HoxA11 (Kim et al., 2003; Gao et al., 2004). As a first step to determine if HoxA11 may regulate *IGFBP1* expression, we explored their expression in previously generated RNA-Seq data human endometrial stromal fibroblasts (ESFs) and ESFs differentiated and decidual stromal cells (DSCs) (Lynch et al., 2015) and found that both were expressed more highly in DSCs than ESFs. Next, we silenced endogenous *HoxA11* expression using siRNA and assayed *IGFBP1* expression 48hrs after transfection and decidualization by quantitative real-time PCR (qRT-PCR). We found that treatment with a *HoxA11* specific siRNA lead to a strong knockdown of *HoxA11* expression and a dramatic up-regulation of *IGFBP1* compared to control siRNA (**Fig. 1B**), suggesting that HoxA11 represses *IGFBP1* expression. To test if HoxA11 represses transcription through the IRS we co-transfected a mouse *HoxA11* expression vector and the luciferase reporter vector pGL2-3xIRS[*luc*/3xIRS], which contains the SV40 promoter and 3 repeats of the IRS, into HeLa cells and assayed luciferase expression 48hrs after transfection; HeLa cells do not express *HoxA11* allowing us measure the effects of HoxA11 on luciferase expression without interference from endogenous *HoxA11*. We found that transfection of the *HoxA11* expression vector repressed luciferase expression ∼64% compared to controls, consistent with HoxA11 being direct a repressor of *IGFBP1* expression (**Fig. 1C**).

Next we mapped the location of the experimentally determined Hox biding-site in the human IRS (Kim et al., 2003; Gao et al., 2004) in relation to the location of H3K27me3 peaks (a marked ‘poised’ *cis*-regulatory elements) identified from human ESFs using ChIP-chip (Grimaldi et al., 2011), as well as binding-sites for co-repressors and DNaseI clusters identified by ChIP-Seq from ESFs (Lynch et al., 2015), to determine if the IRS has the signature of a negative regulatory element for *IGFBP1*. Indeed, the Hox binding site is located within a DNaseI hypersensitivity cluster and an H3K27me3 peak that is lost upon differentiation of ESFs into decidual stromal cells (**Fig. 1D**) and overlaps with ENCODE ChIP-Seq peaks for several co-repressors including YY, SIN3A, and HDAC2 (**Fig. 1D**). Thus we conclude that HoxA11 may directly repress *IGFBP1* expression in association with co-repressors such as SIN3A, YY1, and HDACs.

### The HoxA11 polyalanine tract is a repressor domain

To determine if the HoxA11 polyalanine tract functions in transcriptional regulation, we assayed the ability of mouse HoxA11 amino (N)-terminal deletion mutants to repress luciferase expression from the pGL2-3xIRS[*luc*/3xIRS] reporter vector in transiently transfected HeLa cells. We found that full-length HoxA11 (CDS) as well as N-terminal deletions up to amino acid 170 (Δ170) strongly repressed luciferase expression. In contrast N-terminal deletions that removed the polyalanine tract (Δ222 and Δ242) and an internal deletion of the alanine tract (ΔAla) lost nearly all of their ability to repress luciferase expression (**Fig. 2A**) suggesting the polyalanine tract is necessary for HoxA11 to repress luciferase expression from the pGL2-3xIRS[*luc*/3xIRS] reporter vector.

To test if the HoxA11 polyalanine tract is sufficient to mediate transcriptional repression, we fused it to the GAL4 DNA-binding domain (GAL4-DBD) and tested the ability of the fusion protein to repress luciferase expression from the reporter vector pGL4.35[*luc2P*/9X*GAL4*UAS], which contains the SV40 enhancer/early promoter and nine repeats of the GAL4 Upstream Activator Sequence (UAS); this sequence drives transcription of the luciferase reporter gene *luc2P* in response to binding of fusion proteins containing the GAL4 DNA binding domain. Consistent with the polyalanine tract being a repressor domain, we found that the polyalanine tract GAL4-DBD fusion protein strongly repressed luciferase expression (**Fig. 2A**).

We next inferred whether HoxA11 repressed luciferase expression through steric interference of co-activator binding, or by recruiting a co-repressor complex by treating cells with trichostatin A (TSA), which selectively inhibits mammalian class I and II histone deacetylases. Consistent with HoxA11 recruiting corepressors, we found that TSA treatment effectively abolished luciferase repression by HoxA11 constructs containing the polyalanine stretch and the polyalanine tract GAL4-DBD fusion protein compared to TSA-free controls (**Fig. 2A**). Thus class I or II histone deacetylases (HDACs) are likely required for the HoxA11 polyalanine tract to mediate repression.

### The HoxA11 polyalanine tract interacts with SIN3A PAH domains

A previous study found that the homeodomain of HoxA11 binds a complex including YY1 and HDAC2 to mediate repression (Luke et al., 2006), suggesting that co-factors may mediate repression from the HoxA11 polyalanine tract. To determine if the polyalanine tract mediates physical interactions between HoxA11 and SIN3A or HDAC1, we tested the ability of FLAG-tagged full length HoxA11 and the ΔAla internal deletion mutant to co-immunoprecipitate with HA-tagged HDAC1 and SIN3A. We found that full length HoxA11 interacted with both HDAC1 and SIN3A, however, the ΔAla internal deletion mutant was unable to interact with either SIN3A or HDAC1 (**Fig. 2B**). These results indicate that the polyalanine tract is critical for the interaction of HoxA11 with SIN3A and HDAC1.

SIN3A and its paralog SIN3B contain two highly conserved PAH domains that interact with transcription factors through a ‘wedged helical bundle’ composed of four α-helices (Spronk et al., 2000; Swanson et al., 2004). These helices form a hydrophobic cleft in PAH domains that bind hydrophobic α-helices and the hydrophobic face amphipathic α-helices in their binding partners (Eilers et al., 1999; van Ingen et al., 2003; Anderson et al., 2009), suggesting that the polyalanine tract in HoxA11 may bind to the PAH domains of SIN3A. Consistent with this hypothesis, we found that V5/His-tagged PAH1 and PAH2 domains of SIN3A co-immunoprecipitated with the polyalanine tract GAL4-DBD fusion protein (**Fig. 2C**), but that neither PAH domain co-immunoprecipitated with a GAL4-DBD tagged HoxA11 construct with an internal deletion of the core AAATSAAAVAAAA residues (ΔCoreAla) of the polyalanine tract (**Fig. 2C**). Taken together these results indicate that the HoxA11 polyalanine tract interacts with the PAH domains of SIN3A, likely recruiting a larger co-repressor complex that includes HDAC1.

### The HoxA11 polyalanine repressor domain evolved in mammals

We previously reported that the HoxA11 polyalanine tract evolved in mammals (Chiu et al., 2000; Roth et al., 2005) suggesting that non-mammalian HoxA11 proteins, which have polyalanine tracts less than 6 residues, may not be able to interact with SIN3A to mediate transcriptional repression from the 3xIRS regulatory element. To test this hypothesis, we assayed the ability of chicken, platypus, short-tailed opossum, mouse, and human, as well as reconstructed ancestral Eutherian, Therian, Mammalian, and Amniote HoxA11 proteins to repress luciferase expression from the pGL2-3xIRS[*luc*/3xIRS] reporter vector in transiently transfected HeLa cells. Similar to our observation for mouse HoxA11, extant and reconstructed HoxA11 proteins from other mammals repressed reporter gene expression by 35-81% depending on the species (**Fig. 3A**). The chicken and ancestral Amniote HoxA11 proteins, however, only repressed reporter gene expression by 11-14% (**Fig. 3A**). Thus the ability of HoxA11 to repress reporter gene expression greatly increased in mammals.

**Figure 3.**
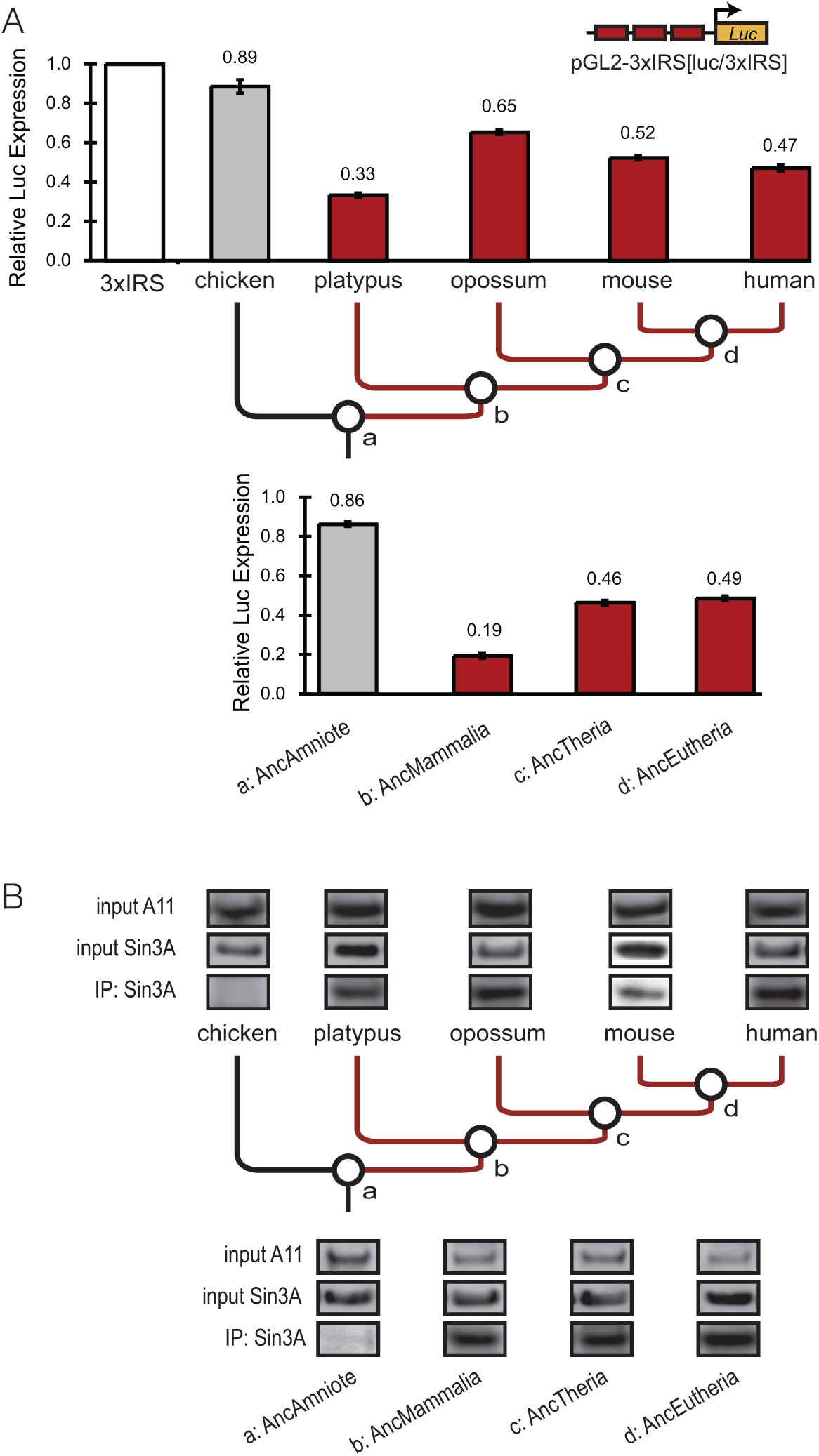
The HoxA11 polyalanine repressor domain evolved in mammals. **(A)** Repression of luciferase expression from the pGL2-3xIRS[*luc*/3xIRS] by extant and reconstructed amniote HoxA11 proteins. Results are shown as fold changes in normalized luciferase (Luc) expression relative to empty vector controls (mean±SD, *n*=4). **(B)** Co-immunoprecipitation of extant (upper) and ancestrally reconstructed (lower) HoxA11 proteins with SIN3A.

Next we tested whether extant and reconstructed ancestral proteins differed in their ability to interact with HA-tagged HDAC1 and SIN3A in co-immunoprecipitation assays. Consistent with our luciferase assay results, V5/His-tagged extant and ancestral HoxA11 proteins from mammals were all able to interact with HDAC1 and SIN3A (**Fig. 3B**). The chicken and ancestral Amniote HoxA11 proteins, however, did not interact with either SIN3A or HDAC1 (**Fig. 3B**). These results demonstrate that the expansion of the polyalanine tract in HoxA11 created a new protein-protein interaction between HoxA11 and SIN3A leading to a new repressor domain in mammals.

### The HoxA11 polyalanine tract evolved in the mammalian stem-lineage

To more precisely determine when the polyalanine track originated, we identified HoxA11 genes from 207 vertebrates including 156 placental mammals, three marsupials, two monotremes, eight sauropsids (including birds, turtles, and squamate reptiles), two species of amphibian, two species of coelacanth, 30 Euteleost fish, and four species of Chondrichthyes. We found that polyalanine tracts longer than six residues were only found in mammals, within mammals polyalanine tracts range from 10-25 residues long (**Fig. 4A**). Parsimony-based ancestral state reconstructions across the HoxA11 genes in our dataset indicates that the common ancestor of amniotes likely had a polyalanine tract that was only 5 residues long, while the common ancestor of mammals had one or two polyalanine tracts that were 12 and 20 residues long, respectively (**Fig. 4B**). While many mammals had contiguous alanine runs 20-22 residues long, we found that only red fox and a ‘long’ allele from domestic dog had repeats longer than 23 residues (**Fig. 4C**). These results indicate that the polyalanine tract expanded in the Mammalian stem-lineage and that purifying selection constrains tract length to be generally less than 23 residues long.

**Figure 4.**
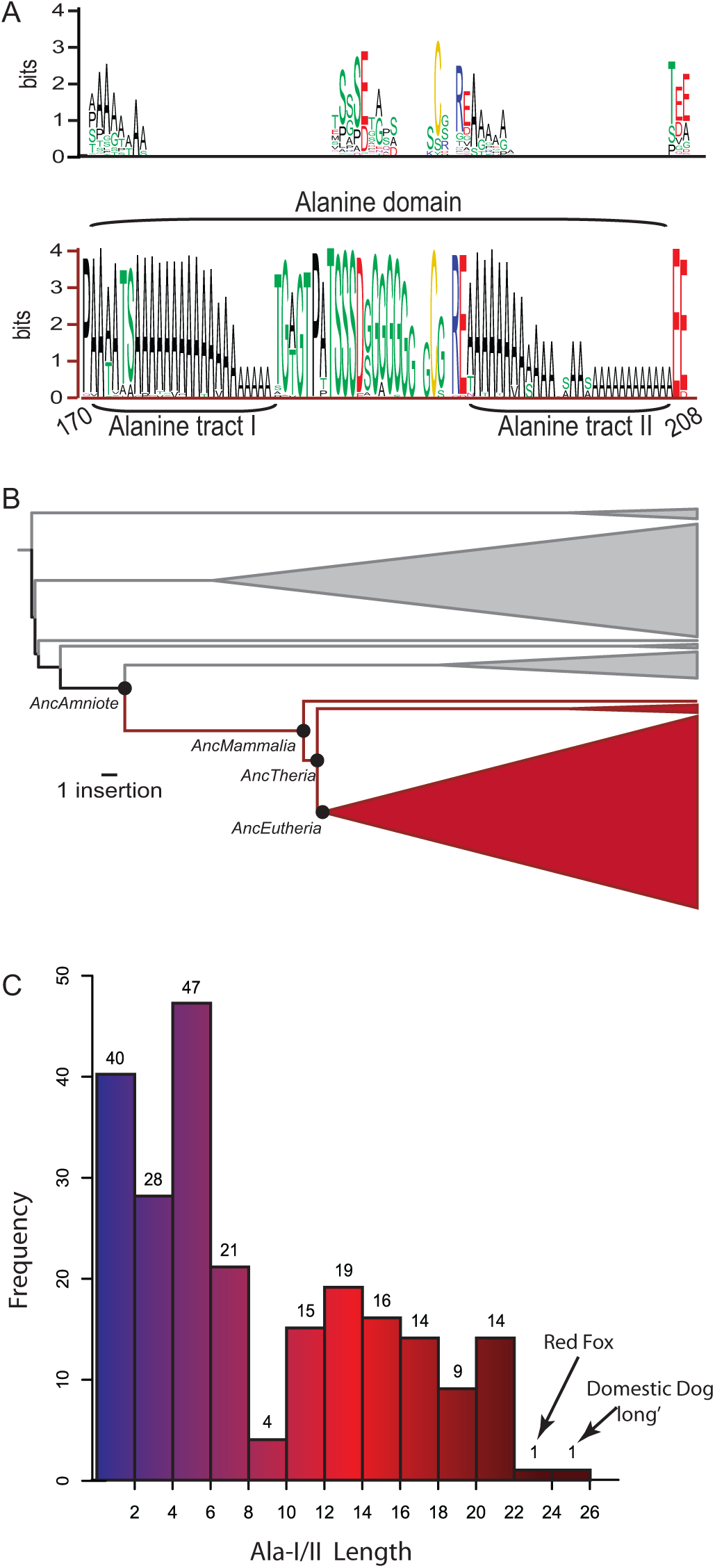
The HoxA11 polyalanine tract evolved in mammals. **(A)** Sequence logo of the polyalanine region from 207 Gnathostomes, including 156 placental mammals, three marsupials, two monotremes, eight sauropsids (birds, turtles, and squamate reptiles), two species of amphibians, two species of coelacanth, 30 Euteleost fish, and four species of Chondrichthyes. Non-mammals upper, mammals lower. Residues are colored according to physicochemical property: Hydrophobic black, positive charge blue, negative charge red, cysteine yellow, and others green. **(B)** Simplified phylogeny of Gnathostomes from which HoxA11 genes were identified. The length of internal branches (black and red) are shown proportional the change in length of the polyalanine domain in that lineage. Mammalian lineages are shown in red, ancestral nodes are also shown. **(C)** Histogram showing the distribution of polyalanine tract lengths across 207 species. Only two species, the longest alleles from red fox and domestic dog, were identified with tracts longer than 23 residues.

### The polyalanine tract may dock into the PAH domains of SIN3

Polyalanine repeats transition from random coil to stable hydrophobic α-helices at ∼10 residues long (Marqusee et al., 1989; Fiori et al., 1993), thus they are biophysically and structurally predisposed to bind the hydrophobic cleft of PAH domains and function as SIN3/HDAC-dependent repressor domains. To better understand the structural basis for the derived interaction between the mammalian HoxA11 polyalanine tract and the PAH domains of SIN3A, we modeled their interaction based on the solution structure of the SIN3A PAH2-HBP1 interaction (Swanson et al., 2004) using Rosetta and RosettaDock (Kim et al., 2004; Lyskov and Gray, 2008). We found that the chicken and ancestral amniote polyalanine tracts were predicted to be mostly unstructured with a short helix (5-8 residues long; **Fig. 5A**), in contrast mammalian polyalanine tracts that were predicted to form long hydrophobic α-helices (**Fig. 5B**). These structural models suggest that the short polyalanine tracts of non-mammalian HoxA11 proteins do not form an α-helix of sufficient length or stability to mediate a functional interaction with PAH domains, whereas the longer tracts found in mammals can form longer amphipathic α-helices that mediate a stable interaction with PAH domains (**Fig. 5C**).

**Figure 5.**
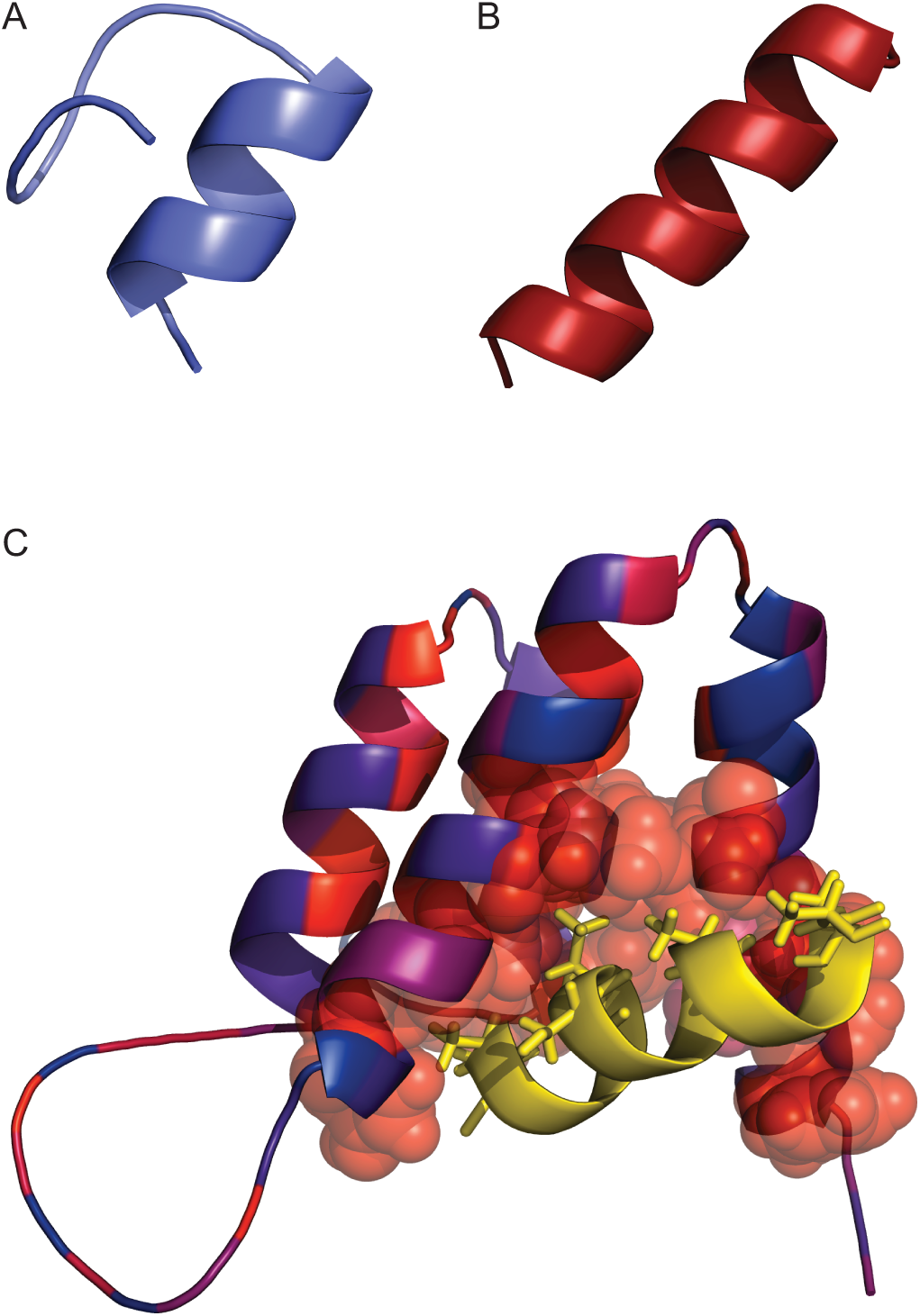
Structural model of the interaction between the HoxA11 polyalanine tract and the PAH domains of SIN3. **(A)** De novo structure model of the polayalanine tract from the ancestral amniote HoxA11 protein. **(B)** De novo structure model of the polayalanine tract from the ancestral Mammalian HoxA11 protein. **(C)** Structural model of the SIN3A PAH2 domain interaction with a segment of the mouse HoxA11 alanine tract (yellow; sequence: AAATSAAAA). Residues in the PAH2 domain are colored according to Kyte-Doolittle hydrophobicity (blue-violet, hydrophilic; red, hydrophobic); residues in the hydrophobic cleft of PAH2 are shown as transparent spheres while residues in the alanine tract predicted to interact with the hydrophobic cleft are shown as sticks.

### A new repressor domain, but at what cost?

While longer polyalanine tracts in mammalian HoxA11 proteins may form more stable interaction interfaces with PAH domains than shorter repeats and thus facilitate their cooption into SIN3-dependant repressor domains, polyalanine tract expansions in transcription factors are associated with numerous human diseases (Goodman et al., 1997; Kjaer et al., 2002; Albrecht et al., 2004; Brown and Brown, 2004). Previous studies, for example, found that polyalanine repeats greater than 23 residues long self-interact to form cytoplasmic aggregates that are excluded from the nucleus (Albrecht et al., 2004; Oma et al., 2007). These results suggest there is a strict upper limit to polyalanine tract length, which is consistent with our observation that only two species possess HoxA11 polyalanine tracts >23 residues long. Given the significant potential for deleterious effects of expanding polyalanine repeats beyond the threshold of ∼23 residues, it is remarkable that selection has maintained long polyalanine repeats in any protein and has coopted at least one into a repressor domain. These observations suggest that while there is a cost associated with functional polyalanine tracts, there is also a benefit for transcription factor function.

## Conclusions

Alanine repeats are common in eukaryotic proteins and emerged during waves of repeat expansions across vertebrates (Albà and Guigo, 2004; Salichs et al., 2009; Haerty and Golding, 2010). Our observations suggest that the predisposition of polyalanine tracts to from stable α-helices may have facilitated their cooption into SIN3-dependant repressor domains independently in many proteins and indicates that amino acid repeats can contribute to the diversification of transcription factor functions (Salichs et al., 2009; Radó-Trilla et al., 2015), despite the potential deleterious consequences of repeat expansions.

## Acknowledgements

This work was funded by a grant from the John Templeton Foundation grants, #12793 and #61329; results presented here do not necessarily reflect the views of the John Templeton Foundation. The funders had no role in study design, data collection and analysis, decision to publish, or preparation of the manuscript. The authors would like to thank A. Rawls (Arizona State University) for providing the SIN3A-HA and HDAC1-HA expression vectors, Y. Shi (Harvard Medical School) for providing the YY1-HA expression vector, and K. H. Correll and J.W. Fondon III (University of Texas Arlington) for providing unpublished HoxA11 polyalanine tracts from vertebrates

## Author contributions

VJL, designed/performed experiments, analyzed data, and wrote the manuscript; GPW wrote the manuscript.

## References

Albà MM, Guigo R. 2004. Comparative Analysis of Amino Acid Repeats in Rodents and Humans. Genome Res 14:549–554.

Albà MM, Santibáñez-Koref MF, Hancock JM. 1999. Amino Acid Reiterations in Yeast Are Overrepresented in Particular Classes of Proteins and Show Evidence of a Slippage-Like Mutational Process. J Mol Evol 49:789–797.

Albrecht AN, Kornak U, B ddrich A, Süring K, Robinson PN, Stiege AC, Lurz R, Stricker S, Wanker EE, Mundlos S. 2004. A molecular pathogenesis for transcription factor associated poly-alanine tract expansions. Hum Mol Genet 13:2351–2359.

Anan K, Yoshida N, Kataoka Y, Sato M, Ichise H, Nasu M, Ueda S. 2007. Morphological Change Caused by Loss of the Taxon-Specific Polyalanine Tract in Hoxd-13. Mol Biol Evol 24:281–287.

Anderson DM, Beres BJ, Wilson-Rawls J, Rawls A. 2009. The homeobox gene Mohawk represses transcription by recruiting the sin3A/HDAC co-repressor complex. Developmental Dynamics 238:572–580.

Brayer KJ, Lynch VJ, Wagner GP. 2011a. Evolution of a derived protein–protein interaction between HoxA11 and Foxo1a in mammals caused by changes in intramolecular regulation. Proc Natl Acad Sci USA 108:E414–E420.

Brayer KJ, Lynch VJ, Wagner GP. 2011b. Evolution of a derived protein-protein interaction between HoxA11 and Foxo1a in mammals caused by changes in intramolecular regulation. Proceedings of the National Academy of Sciences 108:E414.

Briata P, Ilengo C, Van DeWerken R, Corte G. 1997. Mapping of a potent transcriptional repression region of the human homeodomain protein EVX1. FEBS Letters 402:131–135.

Bridgham JT, Ortlund EA, Thornton JW. 2009. An epistatic ratchet constrains the direction of glucocorticoid receptor evolution. Nature 461:515–519.

Brown LY, Brown SA. 2004. Alanine tracts: the expanding story of human illness and trinucleotide repeats. Trends in Genetics 20:51–58.

Chiu CH, Nonaka D, Xue L, Amemiya CT, Wagner GP. 2000. Evolution of Hoxa-11 in lineages phylogenetically positioned along the fin-limb transition. Mol Phylogenet Evol 17:305–316.

Eilers AL, Billin AN, Liu J, Ayer DE. 1999. A 13-Amino Acid Amphipathic alpha Helix Is Required for the Functional Interaction between the Transcriptional Repressor Mad1 and mSin3A. J Biol Chem 274:32750–32756.

Emili A, Greenblatt J, Ingles CJ. 1994. Species-specific interaction of the glutamine-rich activation domains of Sp1 with the TATA box-binding protein. Mol Cell Biol 14:1582–1593.

Faux NG, Bottomley SP, Lesk AM, Irving JA, Morrison JR, la Banda de MG, Whisstock JC. 2005. Functional insights from the distribution and role of homopeptide repeat-containing proteins. Genome Res 15:537–551.

Fiori WR, Miick SM, Millhauser GL. 1993. Increasing sequence length favors alpha-helix over 310-helix in alanine-based peptides: Evidence for a length-dependent structural transition. Biochemistry 32:11957–11962.

Fondon JWI, Garner HR. 2004. Molecular origins of rapid and continuous morphological evolution. Proceedings of the National Academy of Sciences 101:18058–18063.

Galant R, Carroll SB. 2002. Evolution of a transcriptional repression domain in an insect Hox protein. Nature 415:910–913.

Gao J, Mazella J, Tseng L. 2004. Hox Proteins Activate the IGFBP-1 Promoter and Suppress the Function of hPR in Human Endometrial Cells. DNA and Cell Biology 21:819–825.

Gerber HP, Seipel K, Georgiev O, Hofferer M, Hug M, Rusconi S, Schaffner W. 1994., Transcriptional activation modulated by homopolymeric glutamine and proline stretches. Science 263:808–811.

Goodman FR, Mundlos S, Muragaki Y, Donnai D, Giovannucci-Uzielli ML, Lapi E, Majewski F, McGaughran J, McKeown C, Reardon W, Upton J, Winter RM, Olsen BR, Scambler PJ. 1997. Synpolydactyly phenotypes correlate with size of expansions in HOXD13 polyalanine†tract. Proceedings of the National Academy of Sciences 94:7458–7463.

Grimaldi G, Christian M, Steel JH, Henriet P, Poutanen M, Brosens JJ. 2011. Down-Regulation of the Histone Methyltransferase EZH2 Contributes to the Epigenetic Programming of Decidualizing Human Endometrial Stromal Cells. Molecular Endocrinology 25:1892–1903.

Haerty W, Golding GB. 2010. Genome-wide evidence for selection acting on single amino acid repeats. Genome Res.

Heffer A, Shultz JW, Pick L. 2010. Surprising flexibility in a conserved Hox transcription factor over 550 million years of evolution. Proceedings of the National Academy of Sciences 107:18040–18045.

Janody F, Sturny R, Schaeffer VR, Azou Y, Dostatni N. 2001. Two distinct domains of Bicoid mediate its transcriptional downregulation by the Torso pathway. Development 128:2281–2290.

Karlin S, Brocchieri L, Bergman A, Mrazek J, Gentles AJ. 2002. Amino acid runs in eukaryotic proteomes and disease association. Proc Natl Acad Sci USA 99:333–338.

Karlin S, Burge C. 1996. Trinucleotide repeats and long homopeptides in genes and proteins associated with nervous system disease and development. Proceedings of the National Academy of Sciences 93:1560–1565.

Kashi Y, King DG. 2006. Simple sequence repeats as advantageous mutators in evolution. Trends in Genetics 22:253–259.

Kim DE, Chivian D, Baker D. 2004. Protein structure prediction and analysis using the Robetta server. Nucl Acids Res 32:W526–531.

Kim JJ, H S T, Akbas GE, Foucher I, Trembleau A R C J, Fazleabas AT, Unterman TG. 2003. Regulation of Insulin-like Growth Factor Binding Protein-1 promoter activity be FKHR and HOXA10 in primate endometrial cells. Biol Reprod 68:24–30.

King DG, Soller M, Kashi Y. 1997. Evolutionary tuning knobs. Endeavour 21:36–40.

Kjaer KW, Hedeboe J, Bugge M, Hansen C, Friis-Henriksen K, Baeksted M, Niels V, Tommerup N, Opitz J. 2002. HOXD13 polyalanine tract expansion in classical synpolydactyly type Vordingborg. American Journal of Medical Genetics 110:116–121.

Löhr U, Yussa M, Pick L. 2001. Drosophila fushi tarazu: a gene on the border of homeotic function. Curr Biol 11:1403–1412.

Luke MP-S, Sui G, Liu H, Shi Y. 2006. Yin Yang 1 Physically Interacts with Hoxa11 and Represses Hoxa11-dependent Transcription. J Biol Chem 281:33226–33232.

Lynch VJ, May G, Wagner GP. 2011. Regulatory evolution through divergence of a phosphoswitch in the transcription factor CEBPB. Nature 480\:383\–386\.

Lynch VJ, Nnamani MC, Kapusta A, Brayer K, Plaza SL, Mazur EC, Emera D, Sheikh SZ, Grutzner F, Bauersachs S. 2015. Ancient transposable elements transformed the uterine regulatory landscape and transcriptome during the evolution of mammalian pregnancy. Cell Rep 10:551–561.

Lynch VJ, Wagner GP. 2008. Resurrecting the role of transcription factor change in developmental evolution. Evolution 62:2131–2154.

Lyskov S, Gray JJ. 2008. The RosettaDock server for local protein-protein docking. Nucleic Acids Res 36:W233–W238.

Marqusee S, Robbins VH, Baldwin RL. 1989. Unusually stable helix formation in short alanine-based peptides. Proceedings of the National Academy of Sciences 86:5286–5290.

Maurer P, T’Sas F, Coutte L, Callens N, Brenner C, Van Lint C, de Launoit Y, Baert J-L. 2003. FEV acts as a transcriptional repressor through its DNA-binding ETS domain and alanine-rich domain. Oncogene 22:3319–3329.

Nnamani MC, Ganguly S, Erkenbrack EM, Lynch VJ, Mizoue LS, Tong Y, Darling HL, Fuxreiter M, Meiler J, Wagner GP. 2016. A derived allosteric switch underlies the evolution of conditional cooperativity between HOXA11 and FOXO1. Cell Rep 15:2097–2108.

Oma Y, Kino Y, Toriumi K, Sasagawa N, Ishiura S. 2007. Interactions between homopolymeric amino acids (HPAAs). Protein Science 16:2195–2204.

Orr HT, Zoghbi HY. 2007. Trinucleotide Repeat Disorders. Annual Review of Neuroscience 30:575–621.

Ortlund EA, Bridgham JT, Redinbo MR, Thornton JW. 2007. Crystal Structure of an Ancient Protein: Evolution by Conformational Epistasis. Science 317:1544–1548.

Radó-Trilla N, Arató K, Pegueroles C, Raya A, la Luna de S, Albà MM.M 2015. Key Role of Amino Acid Repeat Expansions in the Functional Diversification of Duplicated Transcription Factors. Mol Biol Evol 32:2263–2272.

Ronshaugen M, McGinnis N, Mcginnis W. 2002. Hox protein mutation and macroevolution of the insect body plan. Nature 415:914–917.

Roth JJ, Breitenbach M, Wagner GP. 2005. Represson domain and nuclear localization signal of the murine Hoxa-11 protein are located in the homeodomain: no evidence for role of poly-Alanine stretches in transcriptional repression. J Exp Zool Part B (Mol Dev Evol) 304B:468–475.

Salichs EL, Ledda A, Mularoni L, Alb MM, la Luna de S. 2009. Genome-Wide Analysis of Histidine Repeats Reveals Their Role in the Localization of Human Proteins to the Nuclear Speckles Compartment. PLoS Genet 5:e1000397.

Spronk CAEM, Tessari M, Kaan AM, Jansen JFA, Vermeulen M, Stunnenberg HG, Vuister GW.M 2000. The Mad1-Sin3B interaction involves a novel helical fold. Nat Struct Mol Biol 7\:1100\–1104\.

Swanson KA, Knoepfler PS, Huang K, Kang RS, Cowley SM, Laherty CD, Eisenman RN, Radhakrishnan I. 2004. HBP1 and Mad1 repressors bind the Sin3 corepressor PAH2 domain with opposite helical orientations. Nat Struct Mol Biol 11\:738\–746\.

Thornton JW, Need E, Crews D. 2003. Resurrecting the Ancestral Steroid Receptor: Ancient Origin of Estrogen Signaling. Science 301:1714–1717.

van Ingen H, Lasonder E, Jansen JFA, Kaan AM, Spronk CAEM, Stunnenberg HG, Vuister GW. 2003. Extension of the Binding Motif of the Sin3 Interacting Domain of the Mad Family Proteins. Biochemistry 43:46–54.

Wilson AC. 1975. Evolutionary importance of gene regulation. Stadlar symposia (Uni Missouri), 1975:117–134.

Xiao H, Jeang K-T. 1998. Glutamine-rich Domains Activate Transcription in Yeast Saccharomyces cerevisiae. J Biol Chem 273:22873–22876.

Young ET, Sloan JS, Van Riper K. 2000. Trinucleotide Repeats Are Clustered in Regulatory Genes in Saccharomyces cerevisiae. Genetics 154:1053–1068.

